# Proteolytic Remodeling of 3D Bioprinted Tumor Microenvironments

**DOI:** 10.1101/2023.06.22.546066

**Authors:** Fatemeh Rasti Boroojeni, Sajjad Naeimipour, Philip Lifwergren, Annelie Abrahamsson, Charlotta Dabrosin, Robert Selegård, Daniel Aili

## Abstract

In native tissue, remodeling of the pericellular space is essential for cellular activities and is mediated by tightly regulated proteases. Protease activity is dysregulated in many diseases, including many forms of cancer. Increased proteolytic activity is directly linked to tumor invasion into stroma, metastasis, and angiogenesis as well as all other hallmarks of cancer. Here we show how integrated 3D bioprinted structures with distinctly different responses to proteolytic activity can be utilized for systematic investigation of proteolytic remodeling of the extra cellular matrix and the impact of stromal cells on protease driven processes. Bioprinted structures combining non-degradable and degradable hydrogels were designed and demonstrated to be selectively degraded by proteases allowing for protease-mediated material reorganization with high spatial resolution. Bioprinting of tumor microenvironments combining bioinks with different susceptibilities to proteolytic degradation shows that breast cancer cell proliferation, migration into stromal compartments, and spheroid size are significantly increased in protease degradable hydrogels, but only in the presence of fibroblasts. Proteolytic remodeling of the tumor microenvironment has a significant effect on tumor progression and is drastically influenced by the intimate crosstalk between fibroblast and breast cancer cells.

## Introduction

Proteolytic degradation of the extracellular matrix (ECM) is essential for the dynamic remodeling of the cellular microenvironment ^1, 2^. The remodeling of the ECM influences cell differentiation, generation, and maintenance of stem cell niches, branching morphogenesis, angiogenesis, bone remodeling, wound healing, and tissue morphogenesis during embryonic development ^3, 4^. A deregulated proteolytic activity profile result in congenital defects and pathological processes including tissue fibrosis and cancer ^5^. In cancer, proteases secreted by cancer cells, cancer associated fibroblasts (CAFs), and other non-malignant host cells in the tumor microenvironment (TME) are shaping the TME and contributing to the progression of the disease ^6, 7^. Engineering of biologically relevant tissue and disease models thus requires ECM mimicking materials that can respond in an appropriate way to proteolytic activity. Protease responsive hydrogels incorporating protease cleavable biopolymers or cross-linkers ^8, 9^ have been widely explored for three-dimensional (3D) cell culture of cancer cells and have also been explored as components in bioinks for 3D bioprinting ^10, 11^. 3D bioprinting has emerged as a promising technology for the fabrication of more sophisticated tumor models ^12, 13^. Extrusion based bioprinting ^14^ relies on layer-by-layer deposition of cell laden biomaterials ^15^. The biomaterial is typically a hydrogel or a hydrogel precursor ^16^, which provides a hydrated ECM mimicking environment ^17, 18^ that both act as a scaffold and ideally protects the cells from detrimental shear forces during printing.

Protein or peptide-based hydrogels, including hydrogels comprised of e.g., collagen, gelatin, and elastin, are inherently degradable by proteases. Hydrogels prepared using synthetic polymers, such as poly(ethylene glycol), or carbohydrates, such as hyaluronan or alginate, are typically not protease responsive but can be cross-linked using proteins or peptides targeting specific groups of proteases ^8, 19–22^. Because of their central role in ECM turnover in both healthy tissue and disease, numerous peptides cleaved by specific types or groups of matrix metalloproteinases (MMPs) and other proteases, have been identified and are widely used as components in protease-responsive hydrogels. Matrix metalloproteinases (MMPs) are a family of zinc-dependent endopeptidases that play central roles in numerous physiological processes, such as organ development and tissue remodeling ^1^. In healthy tissue, the expression of MMPs is tightly regulated, whereas in almost every type of human cancer, their expression is significantly upregulated ^23, 24^. The complex pattern of MMP over-expression in the TME depends on both the cancer cells and the tumor stromal cells, and results in proteolytic cleavage of extracellular matrix (ECM) components, growth-factor precursors and growth-factor binding proteins, cell adhesion molecules, receptor tyrosine kinases, and other proteinases ^24^. Fibroblasts are the major cellular component of the TME and central for ECM deposition and remodeling. The degradation of the ECM promotes cancer cell invasion and metastasis, which can be further enhanced by the proteolytic activation of e.g. transforming growth factor β (TGF-β). MMPs are also involved in tumor angiogenesis by regulating the bioavailability of vasculature endothelial growth factor (VGEF). However, the complex crosstalk between different cell types in the TME and the lack of defined 3D cancer models make systematic studies of the role and mechanisms of proteolytic remodeling very challenging.

Here we show a well-defined protease responsive hydrogel-system for 3D bioprinting of breast cancer models that delineates the role of proteases in tumor progression. We designed a set of hyaluronan (HA) based extracellular matrix mimicking (ECM) hydrogels using peptide and poly(ethylene glycol) (PEG) based cross-linkers. Whereas the peptide cross-linked hydrogels where rapidly and efficiently proteolytically degraded, hydrogels cross-linked by PEG remained intact, enabling bioprinting of spatially defined structures that responded differently to proteolytic activity. Bioprinted mimics of the tumor microenvironment comprising both human breast cancer cells and primary fibroblasts in separate but adjacent compartments were generated and allowed for a systematic exploration of the effect of proteolytic ECM remodeling on cancer cell proliferation migration, and spheroid growth. A distinct and synergistic relationship between hydrogel degradability and cancer cell growth, fueled by the presence of fibroblasts, was observed.

Bioprinting facilitates the organization of multiple cell types and different ECM-mimicking materials in spatially defined 3D architectures, providing significantly better possibilities to mimic the spatial heterogeneity and complexity of the TME. Our findings show that proteolytic remodeling of the ECM accelerates tumor progression and occurs as concerted interactions between cancer cells and fibroblasts, which can pave the way for development of new therapeutic strategies and more elaborate tumor models.

### Experimental section

Unless otherwise specified, all chemicals were obtained from Merck Life Science AB (Stockholm, Sweden) and used as obtained.

### HA-BCN synthesis

HA-BCN was synthesized as previously described ^25^. Briefly, 500 mg of sodium hyaluronate (100-150 kDa, Lifecore Biomedical Inc., Chaska, USA) was dissolved in 40 ml of 4-morpholineethanesulfonic acid (MES buffer, 100 mM, pH 7). In a separate container, 100 mg of *N*-[(1*R*,8*S*,9*s*)-Bicyclo[6.1.0]non-4-yn-9-ylmethyloxycarbonyl]-1,8-diamino-3,6-dioxaoctane (BCN-NH_2_) was dissolved in 6 ml of a 5:1 acetonitrile: water solution, followed by addition of 83 mg of 1-hydroxybenzotriazole (HOBt) and 236 mg of N-(3-Dimethylaminopropyl)-N′-ethylcarbodiimide hydrochloride (EDC), and subsequently added to the hyaluronic acid solution and allowed to react at room temperature for 24 hours. The solution was dialyzed (molecular weight cutoff 6-8 kDa, Spectra/Por RC, Spectrum Laboratories, New Brunswick, USA) against 10% acetonitrile for 1 day, followed by changing the dialysis solution to MQ-water (18.2 MΩ cm^-1^) and continued the dialysis for 5 days. The degree of substitution was determined by using proton nuclear magnetic resonance (^1^H-NMR 500Hz, Oxford Instruments, Abingdon, United Kingdom) in D_2_O.

### Peptide design and synthesis

The cyclic RGD peptide (cRGD-Az) with an azide moiety and the sequence c(RGDfK(Az)) was synthesized as described earlier ^26^. The VPM peptide (VPM-Az_2_) with the sequence Ac-(K(Az)-GRDVPMSMRGGDR-K(Az))-CONH_2_ was synthesized on a microwave assisted automated peptide synthesizer (Liberty Blue, CEM, Matthews, NC, USA) using standard fluroenylmethoxycarbonyl (Fmoc) chemistry in a 250 μmol scale. The first amino acid Fmoc-Lys(Az)-OH (Iris Biotech GmbH, Marktredwitz, Germany) was coupled to a rink amide resin (ProTide, LL, loading 0.19 mmol/g) using a five-fold excess of amino acid, *N*,*N*’-Diisopropylcarbodiimide (DIC) and a ten-fold excess of Oxyma Pure as base. The reaction mixture was heated to 90 °C using microwave conditions for 2 min. The coupling procedure was performed twice. Subsequent deprotection of the Fmoc protection group was accomplished by treatment with 20% piperidine in DMF (v/v) at 90 °C for 1 min. The remaining amino acids were attached using the same procedure. N-terminal acetylation after final Fmoc deprotection was achieved by treating the peptide with 50% acetic anhydride in DMF (v/v) for 1 hour. The crude peptide was obtained by cleavage and global deprotection using trifluoroacetic acid (TFA), water, and triisopropylsilane (TIS) (95:2,5:2,5, v/v/v) for 3 hours before being filtered, concentrated, and precipitated twice in ice-cold diethyl ether. The peptides were purified using a reversed phase C-18 column (ReproSil Gold) attached to a HPLC system (Dionex Ultimate 3000 LC, Thermo Scientific^TM^, Thermo Fisher Scientific Waltham, Massachusetts, USA). After concentrating and freeze drying, the purity of the peptide was confirmed by both HPLC and MALDI-ToF mass spectrometry (UltrafleXtreme, Bruker Daltonics, Billerica, MA, USA).

### Hydrogel synthesis and analysis

All hydrogels were prepared by dissolving the HA-BCN and the cross-linker separately. First, HA-BCN was dissolved at a concentration of 20 mg ml^-1^ in 140 mM sodium chloride, 2.7 mM potassium chloride and 10 mM phosphate, pH 7.4 (PBS). Xanthan gum (XG) powder was added to the HA-BCN solution (20 mg ml^-1^) and carefully mixed to achieve concentrations of 0.5%, 1%, and 1.5% (w/v) XG. The different solutions were referred to as HA, HA/0.5% XG, HA/1% XG, and HA/1.5% XG. For protease-degradable and non-degradable hydrogels, two different cross-linkers, protease-degradable peptide VPM-Az_2_ and non-degradable PEG-bis-azide (Mw = 1100 Da, PEG-Az_2_) were dissolved in PBS at concentrations of 20 and 12.34 mg ml^-1^, respectively. The hydrogels were obtained by mixing HA-BCN ± XG and cross-linker solutions at 7:1 volume ratio. The hydrogels were also supplemented with 100 µM cRGD-Az peptide for all conditions. Oscillatory rheology was conducted using a Discovery HR-2 rheometer (TA Instruments, New Castle, USA). Bioink viscosity was measured at a shear rate of 0.01 to 100 s^-1^. The shear recovery test was conducted using oscillatory time sweeps at 1 Hz and changing from high (250%) to low (0.5%) strain every two minutes. The cross-linking kinetics of the hydrogels was monitored using a 20 mm diameter and 1° cone-plate geometry at a frequency of 1 Hz and a strain of 1 % at 37 °C directly after depositing the samples (50 µl) on the rheometer stage and immediately after mixing the HA-BCN (±XG) and the cross-linkers, followed by frequency and amplitude sweeps. To measure the rheological properties of hydrogels after swelling, 30 µl of the hydrogel was poured between two glass slides directly after mixing the components and covered by parafilm separated by a 500 µm spacer. The hydrogels were allowed to cross-link for 2 hr at 37°C and then immersed in PBS and equilibrated for 24 hr. The rheological characterization of the obtained hydrogel disks was performed using an 8 mm parallel plate geometry with a gap high of ∼500 µm. All rheological measurements were performed in a linear viscoelastic region as confirmed by amplitude sweep.

### Degradation of hydrogels by collagenase

To study the proteolytic degradation of the hydrogels, HA-BCN was labeled with sulfo-Cy5-Az. Briefly, 10 µM sulfo-Cy5-Az (Lumiprobe GmbH, Hanover, Germany) was allowed to react with 20 mg ml^-1^ HA-BCN for 12 hr followed by dialysis against MQ-water and then lyophilized. The resulting Cy5-labeled-HA-BCN was further utilized to track the degradation of hydrogels when exposed to different concentrations of collagenase (0.005, 0.05, and 0.5 mg ml^-1^). After mixing the hydrogel components, 20 µl of the final solution was cast at the bottom of a 1 ml plastic cuvette followed by centrifugation of the cuvette. After cross-linking for 2 hr at 37 °C, 1 ml of 30 mM 4-(2-hydroxyethyl)-1-piperazineethanesulfonic acid (HEPES buffer, pH 7) supplemented with 5 mM Ca^2+^ was added and allowed to equilibrate for 2 hr at 37 °C. Hydrogel degradation was recorded by reading the fluorescence emission at 620 nm before and after addition of collagenase type 1 (Col-1) at different concentrations (0.005, 0.05, and 0.5 mg ml^-1^ in 30 mM HEPES buffer pH 7, supplemented with 5 mM Ca^2+^) at regular time intervals for 48 hr. After the final measurement, 50 µl of 10 mg ml^-1^ of hyaluronidase from bovine testes (Type 1-S, 400-1000 units/mg solid) was added to each sample followed by 24 hours incubation for 2 hr at 37 °C to fully disassemble any remaining parts of the hydrogel to determine the percentage of the degraded hydrogel. For collagenase-mediated remodeling of bioprinted structures, bioprinted structures were prepared using the different hydrogel compositions and then immersed in 1 mg ml^-1^ Col-1 in 30 mM HEPES pH 7 with 5 mM Ca^2+^ for 24 hr. The structures were imaged every 2 hours.

### Cells and 3D cell culture

Primary human dermal fibroblasts were obtained from skin biopsies from healthy patients. All experiments involving human tissue and cells were performed under ethical approval from the Swedish Ethical Review Authority (no. 2018/97-31) and in accordance with ethical standards postulated by Linköping University and Swedish and European regulations. Skin was obtained from healthy female patients undergoing routine breast reduction surgery and de-identified. Fibroblasts were isolated and expanded using standard sterile protocols ^27^. Briefly, skin samples were repeatedly washed in sterile PBS and subcutaneous fat and epidermis were mechanically removed. The remaining dermis was cut into 1 × 3 mm^2^ pieces and placed in Dulbecco’s modified Eagle’s medium (DMEM, Gibco Thermo Fisher Scientific, Paisley, UK) with 165 U ml^-1^ collagenase (Gibco Thermo Fisher Scientific, Paisley, UK) and 2.5 mg ml^-1^ dispase (Gibco Thermo Fisher Scientific, Paisley, UK) and incubated at 37 °C, 5% CO_2_, and 95% humidity overnight. The pooled supernatants were centrifuged for 5 min at 365 × g and the resulting cell pellet was re-suspended in fibroblast medium (DMEM supplemented with 10% Fetal Calf Serum, 50 U/ml Penicillin and 50 mg/ml Streptomycin). Cells were seeded into 75 cm^2^ culture flasks (Falcon, Corning Inc; Corning, NY, USA) with fibroblast-medium and kept in an incubator at 37 °C, 5% CO_2_, and 95% humidity. The medium was changed three times per week. The isolated fibroblasts were cultured in high glucose DMEM medium supplemented with 10% FBS and 1% penicillin-streptomycin. Medium was changed every 3 days, and cells were passaged at 70% confluency. Fibroblasts used in this study were in passages 5-8. For cell encapsulation and bioprinting, fibroblasts were lifted from tissue culture flask with 0.25 % trypsin-EDTA solution for 10-15 min at 37°C, followed by counting, centrifuging, pelleting, and resuspending in high glucose DMEM medium supplemented with 10% FBS and 1% penicillin-streptomycin. To inactivate all trypsin in cell suspension the previous steps were repeated in FBS. To achieve a final cell density of 1 × 10^6^ cells ml^-1^ and 3 × 10^6^ cells ml^-1^ in hydrogel for 3D cell culture and bioprinting, respectively, the desired number of fibroblasts were pelleted and resuspended in the mixture of cross-linker and cRGD-Az. The cell-laden hydrogels were prepared by mixing the cell suspension containing cells, cross-linker (VPM-Az_2_ or PEG-Az_2_), and cRGD-Az with HA-BCN ± XG in culture medium at the ratio described above. The hydrogels were cross-linked for 2 hr in the incubator at 37 °C followed by the addition of culture medium to each sample. The culture medium was changed every 2-3 days. MCF-7 cells were a kind gift from Prof. Charlotta Dabrosin (Division of Surgery, Orthopedics and Oncology (KOO), Linköping University). MCF-7 were cultured in DMEM medium supplemented with 10% FBS and 1 % penicillin-streptomycin. The culture medium was changed every 2-3 days, and cells were split at 70% confluency. For bioprinting, MCF-7 cells were lifted from tissue culture flask using TrypLE^TM^ solution for 5 min at 37°C and resuspended in fresh culture medium to inactivate TrypLE^TM^. The concentration of cells in the suspension was determined using trypan blue before plating. Subsequently, the desired number of cells were resuspended in a solution of cross-linker and cRGD-Az and then mixed with HA-BCN ± XG and immediately used for bioprinting.

### 3D Bioprinting

Bioprinting was performed using a BIO X^TM^ 3D bioprinter (Cellink AB, Gothenburg, Sweden). All bioprinting was conducted using a standard pneumatic printhead loaded with a 3 ml syringe attached to a 25-gauge needle. The viability of cells (fibroblasts) in the printed hydrogels was assessed by LIVE/DEAD staining (Biotium, Fremont, California, USA) 24 hr after printing. Briefly, 50 µl of hydrogels containing 3 × 10^6^ cells ml^-1^ were extruded, and the printed structures (n=3) were allowed to cross-link at 37 °C for 2 hr followed by the addition of culture medium and incubation at 37 °C overnight. In addition, to compare the viability of cells in printed and pipetted hydrogels, 50 µl of hydrogels were also pipetted in a 24-well plate and the aforementioned steps were repeated for this group of hydrogels. After 24 hr the structures were washed with PBS and immersed into the LIVE/DEAD staining solution (2 µM calcein AM and 4 µM ethidium homodimer-1 in PBS) for 30 min at room temperature. To quantify the viability of cells, a stack with 30 slices (∼220 µm in total) was imaged using 10 X objective on a confocal fluorescent microscope (LSM700, Zeiss, Oberkochen, Baden-Württemberg, Germany) and converted to a 2D projection. Heart-shaped structures with or without MCF-7 (4 × 10^6^ cells ml^-1^) were printed in a 6-well-plate using bioinks labeled with sulfo-Cy5-Az. After 2 hr incubation at 37 °C, the structures were covered by bioink with or without fibroblasts (3 × 10^6^ cells ml^-1^) followed by 2 hr incubation at 37 °C. The printed structures were subsequently immersed in culture medium and placed in the incubator at 37 °C and 5% CO_2_. For all conditions, the culture medium was changed every 3 days during 14 days of culture.

### Analysis of cell morphology

Encapsulated cells were fixed in 4 % paraformaldehyde in PBS for 30 min and permeabilized with 0.5 % Triton X-100 in PBS for 30 min. The samples were washed thoroughly three times in PBS to remove any remaining components. Samples were incubated in 0.165 µM phalloidin (Phalloidin, CF488A conjugate, green (490/515 nm)) and 2 µg ml^-1^ Hoechst 33258 (355/465 nm) to stain F-actin and nucleus, respectively. After 2 hours, the samples were washed with BPS and used for confocal fluorescent microscopy. A ≈ 400-500 µm stack with 50 slices was performed at 10 X objective on a confocal fluorescent microscope (LSM700). Huygens Professional (SVI, Hilversum, Netherlands) was used for deconvolution and creating rendered surface images as well as the analysis of spheroid volume. To measure the cell area and circularity, the original 3D-stack image was converted to a 2D-projected image using ImageJ software (FIJI)^28^. The cell area and circularity were measured using 2D-projected images with ImageJ software (FIJI). The circularity is defined as 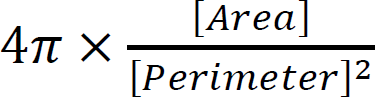.

### Statistical analysis

All experiments were performed in triplicate unless otherwise stated. All data were shown as mean ± standard deviation (SD). Statistical analysis was performed using GraphPad Prism 9 software (San Diego, CA, USA). The specific data analysis used for each data set is mentioned in the figure captions.

## Results and discussion

### Hydrogels and bioinks

Hyaluronan (HA) hydrogels were prepared by cross-linking HA modified with bicyclo[6.1.0]nonyne (BCN) by strain-promoted azide-alkyne cycloaddition (SPAAC) using azide-functionalized cross-linkers (Fig. S1, Supporting Information). To allow for comparison and seamless integration of non-degradable and protease-degradable HA hydrogel structures, two different bifunctional azide (Az) cross-linkers were used (Fig. 1a). As a degradable cross-linker we synthesized a peptide with the sequence K(Az)-GRDVPMSMRGGDR-K(Az) (VPM-Az_2_) (Fig. S2, Supporting Information). As a non-degradable cross-linker, a bifunctional PEG-bis-azide (PEG-Az_2_) was used. VPM-peptides have been widely employed as protease degradable components in hydrogels and is hydrolyzed by e.g., MMP-1 and MMP-2 ^8, 20, 29–31^. To promote cell-hydrogel interactions we further modified the HA-backbone with Az-containing cyclic RGD peptides (cRGD-Az). The molecular weight of PEG-Az_2_ (1100 Da) was chosen to match the size of VPM-Az_2_ to facilitate the fabrication of non-protease degradable (HA-PEG) and protease degradable (HA-VPM) hydrogels with similar mechanical properties and network topologies. The mechanical properties of the hydrogels were found to be highly dependent on the ratio of HA-BCN and cross-linkers. After careful optimization (Table S1, Supporting Information), three ratios of HA-BCN and cross-linkers were selected for further rheological analysis, where a HA-BCN volume ratio of 7:1 showed the fastest gelation and generated hydrogels with suitable stiffness for soft tissue biofabrication (Fig. S3, Supporting Information). This ratio was used in all further experiments. At this ratio, PEG-Az_2_ and VPM-Az_2_ showed similar cross-linking kinetics (Fig. 1b,c). Increasing the amount of cross-linker resulted in faster gelation but softer hydrogels because of the saturation of the BCN groups which prevented further network formation (Fig. S3, Supporting Information).

**Figure 1.**
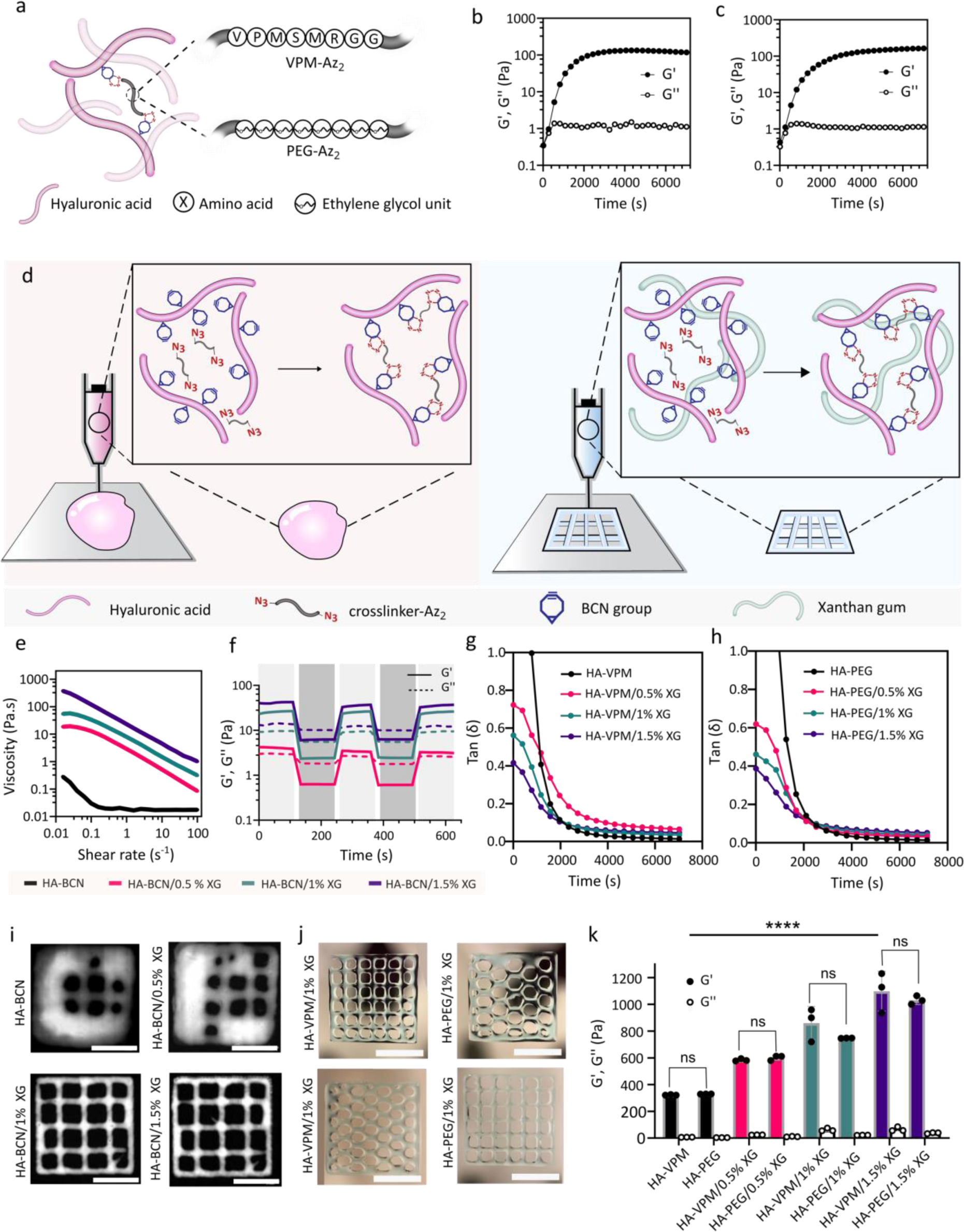
Bioink optimization. **a.** Schematic illustration of hydrogel cross-linking. **b.** Cross-linking kinetics of HA-BCN after addition of VPM-Az_2_. **c.** Cross-linking kinetics of HA-BCN after addition of PEG-Az_2_. **d.** Schematic illustration of hydrogel printing without and with XG. **e**. Continuous flow test of four different bioink compositions using VPM-Az_2_ as cross-linker and increasing concentration of XG, 0.01 – 100 s^-1^ shear rate. **f.** Recovery tests of bioinks prior to cross-linking were conducted by applying cyclic strain of 0.5% (light gray) and 250% (dark gray) at 1 Hz. **g**. Time sweep test showing loss tan(δ) of hydrogels cross-linked with VPM-Az_2._ **h**. Time sweep test showing loss tan(δ) of hydrogels cross-linked with PEG-Az_2_ **i.** 3D printing of grid structures showing increasing shape fidelity upon addition of XG. Scale bar: 5 mm **j.** 3D printing of structures with different infill patterns. Scale bar: 5 mm **k.** Storage (G’) and loss (G“) modulus of HA-VPM and HA-PEG hydrogels ± XG (0.5-1.5%) after swelling in PBS (****P < 0.0001, ns no significant, one-way ANOVA followed by Tukey’s multiple comparisons test, n=3).

The SPAAC cross-linking reaction is biorthogonal and has previously been demonstrated to enable encapsulation and culture of cells with high viabilities ^32, 33^. However, we noticed that the relatively low viscosity of the HA-VPM and HA-PEG hydrogels prior to and during early-stage cross-linking could complicate 3D bioprinting, resulting in poor printing resolution and shape fidelity. Increasing the concentration of HA-BCN can increase the viscosity but narrows the printing window. In addition, higher concentration of HA-BCN results in hydrogels with a final stiffnesses outside the desired range ^34^. To improve printability, we instead added xanthan gum (XG) to the HA-BCN solution prior addition of cross-linkers (Fig. 1d). XG is well tolerated by cells and widely used in bioinks to modulate their viscosity ^35–38^. The effect of increasing XG concentration from 0 to 1.5 % (w/v) were investigated. Addition of 1.5 % (w/v) XG resulted in an increase in viscosity from 450 mPa.s to 275 Pa.s a shear rate of 0.01 s^-1^ (Fig. 1e) ^39^. When increasing the shear rate to 100 s^-1^ to simulate the conditions during bioprinting, we observed a clear shear thinning behavior of the XG-containing hydrogels. In addition, a recovery test was performed at a frequency of 1 Hz by alternating the strain from 0.5 % (low) to 250 % (high) (Fig. 1f). The results showed a higher loss modulus than storage modulus (G“ > G’) at a strain of 250 % indicating a fluid-like behavior, which is desirable for bioprinting. Decreasing the strain to 0.5 % resulted in the opposite, with higher storage modulus than loss modulus (G“ < G’), indicative of a solid-like materials. The shear thinning behavior and the rapid recovery of the hydrogels when going from high to low strain allows for the material to flow during printing but stabilizes the shape of structures after printing, which make the XG-containing hydrogels excellent candidates for extrusion bioprinting. The impact of XG on the printing performance was further studied by monitoring the loss tangent of the hydrogels as a function of XG content (Fig. 1g,h). No major difference in loss tangents were seen for fully cross-linked hydrogels, with and without XG. However, hydrogels without XG showed very high loss tangent during the initial phase of cross-linking, which can result in poor shape fidelity of printed structures. Increasing the concentration of XG drastically decreased the loss tangent during the initial phase of cross-linking, which on the other hand, can facilitate printing ^40, 41^. The impact of XG on printability was further evaluated using different printing parameters (Table S2, Supporting Information) which confirmed that XG concentrations of 1-1.5 % (w/v) drastically improved the printability of the hydrogels (Fig. 1i,j). Without XG, the structures showed very poor shape fidelity and fusion of printed filaments.

In addition, the rheological properties of the hydrogels after gelation and swelling in PBS were measured. As expected, increasing the concentration of XG resulted in stiffer hydrogels irrespectively of which cross-linker that was used (Fig. 1k). Since the type of cross-linker did not significantly affect the rheological properties of the hydrogels we concluded that the HA-PEG and HA-VPM hydrogels can be used interchangeably for controlling and tuning proteolytic degradation of bioprinted structures.

### Cell viability

A bioink must be able to maintain high cell viability, proliferation, and promote the desired phenotypes ^42^. During extrusion-based bioprinting cells can experience significant shear stress that has a negative impact on their viability. The stress depends on the viscosity of the bioink, needle gauge, and the pressure applied to extrude the bioink.^39^ The impact of printing process of bioinks with different concentrations of XG (0, 0.5%, 1%, 1.5%) and the two types of cross-linkers (PEG-Az_2_ and VPM-Az_2_) was evaluated. We observed minimal impact by the printing process on the viability of the cells 24 hours post bioprinting, with no decrease in cell viability for XG concentrations up to 1% (Fig. 2a,b). A slight decrease in cell viability to 89 ± 2% was seen for bioinks containing the highest amount of XG (1.5%), likely due to the higher pressure required for printing these hydrogels. The bioink composition containing 1% XG (HA-VPM/1% XG and HA-PEG/1% XG) showed both high cell viability post bioprinting and good printability and was thus used for the remaining part of the work. The effect of type of cross-linkers (PEG-Az_2_ and VPM-Az_2_) on fibroblast morphology and phenotype was further evaluated (Fig. 2c-f). Both cell density and cell area increased over a period of 14 days, but no differences were seen between the two types of cross-linkers, indicating adequate cell proliferation and cell-material interactions in both cases. Fibroblasts cultured in VPM-Az_2_ cross-linked hydrogels showed a more elongated morphology compared to when cultured in the PEG-Az_2_ cross-linked hydrogels, presumably due to proteolytic hydrogel remodeling in the former case.

**Figure 2.**
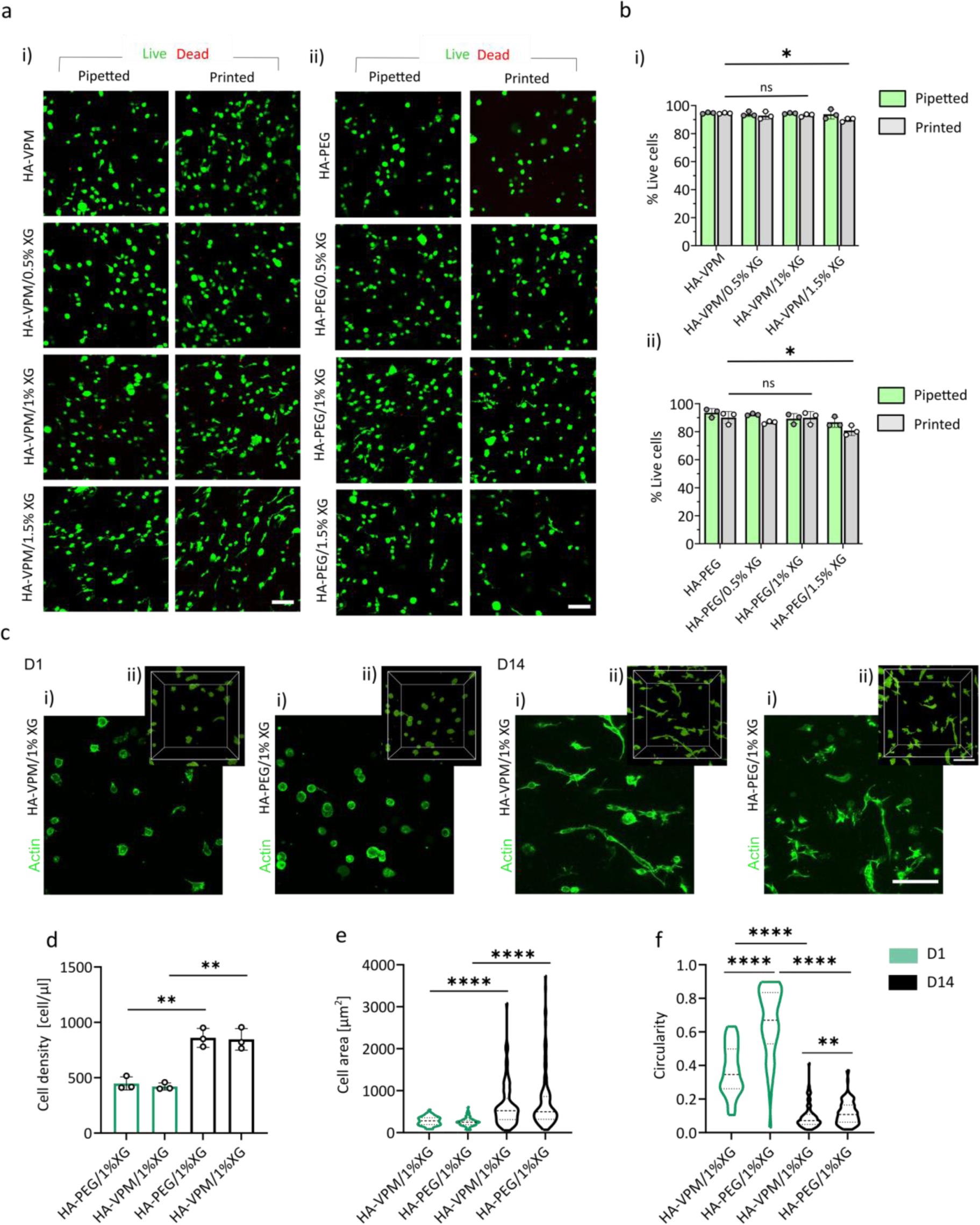
Effect of hydrogel composition on human primary fibroblast viability and morphology. **a.** Live/Dead staining of fibroblasts (green: calcein-AM, red: ethidium homodimer-1) in hydrogel cross-linked with VPM-Az_2_ (i) and PEG-Az_2_ (ii) after 24 hours. Scale bar, 100 µm. **b.** Quantification of fibroblasts viability 24 hr after bioprinting or gentle extrusion through a pipette using different hydrogel compositions cross-linked with (i) VPM-Az_2_ and (ii) PEG-Az_2_ (*, P < 0.0332; ns no significant, two-way ANOVA followed by Sidak’s multiple comparisons test, n=3). **c.** Confocal (i) and rendered surface images (ii) of fibroblasts (green, phalloidin stained actin) cultured in hydrogels cross-linked with VPM-Az_2_ and PEG-Az_2_ at day 1 (D1) and day 14 (D14). Scale bar, 100 µm. **d.** Quantification of fibroblasts cell density (**, P < 0.0021, unpaired t test, n=3). **e.** Quantification of fibroblasts cell area (****, P < 0.0001, Mann-Whitney test). **f**. Circularity of the fibroblasts (****, P < 0.0001, Mann-Whitney test).

### Hydrogel degradation kinetics

The use of the two different cross-linkers allows for tuning of the proteolytic degradation of the hydrogels (Fig. 3a). The rate of degradation was investigated by labeling the HA backbone with a fluorophore (Cy5-Az) and observing the release of HA from the hydrogels after addition of collagenase type I (Col-1). Minor mass loss was seen for the HA-PEG hydrogel, both in the absence and presence of Col-1, indicating release of some non-covalently bound (entangled) HA (Fig. 3b). In contrast, the HA-VPM hydrogels were almost completely degraded after 2 hours at the two highest Col-1 concentrations used (0.05 and 0.5 mg ml^-1^) (Fig. 3c). The rate of degradation decreased with increasing XG concentration (Fig. 3d-g), likely due to the slower diffusion and release of the degradation products rather than slower proteolysis ^43^.

**Figure 3.**
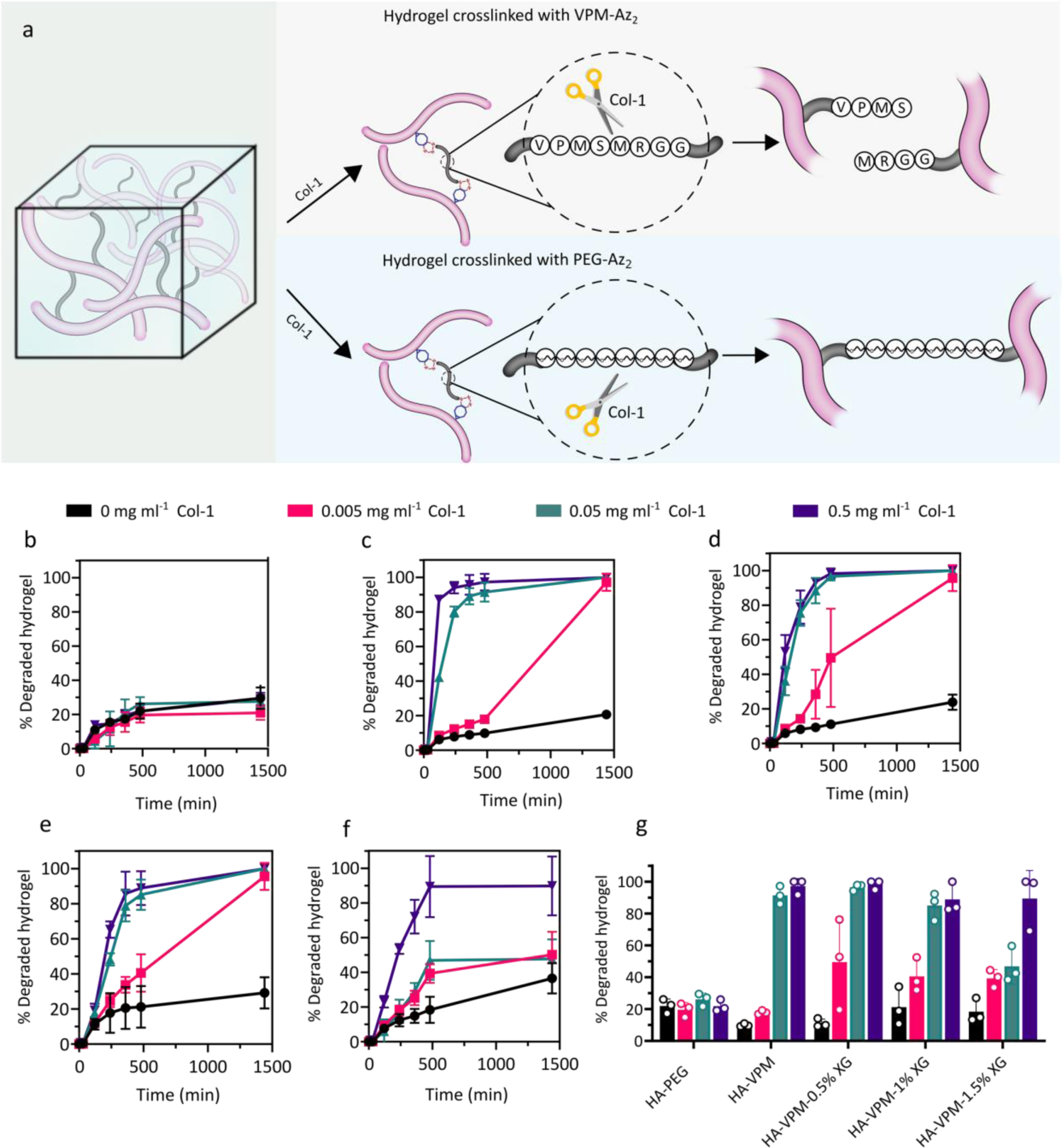
Proteolytic degradation of hydrogels by Col-1. **a.** Schematic illustration of the effect of Col-1 on hydrogels crosslinked by VPM-Az_2_ and PEG-Az_2_. **b.** The degradation rate of HA-PEG hydrogels exposed to 0-0.5 mg ml^-1^ Col-1. **c.** The degradation rate of HA-VPM hydrogel, exposed to 0-0.5 mg ml^-1^ Col-1. **d.** The degradation rate of HA-VPM/0.5% XG hydrogel, exposed to 0-0.5 mg ml^-1^ Col-1. **e.** The degradation rate of HA-VPM/1% XG hydrogel, exposed to 0-0.5 mg ml^-1^ Col-1. **f.** The degradation rate of HA-VPM/1.5% XG hydrogel, exposed to 0-0.5 mg ml^-1^ Col-1. **g.** Degradation of hydrogels after 8 hr incubation with 0-0.5 mg ml^-1^ Col-1.

### Protease remodeling of 3D printed hydrogels

The possibility to seamlessly combine bioinks that respond differently to proteolytic activity was investigated as a new means for remodeling bioprinted structures, including reducing cross-linking density and thus stiffness and swelling and ultimately for selective dissolution of specific printed features. To explore the latter, we printed defined structures utilizing either HA-VPM/1 % XG or HA-PEG/1% XG with a second homogenous printed on top comprising any of the two different hydrogel compositions (Fig. 4a). The final structures were allowed to be fully cross-linked and were then subsequently immersed in a buffer containing Col-1 (0.5 mg ml^-1^) and imaged every 2 hours for 8 hours. The remodeling of the structures in response to Col-1 was highly dependent on the combination of hydrogels (Fig. 4b). The structures printed only with HA-VPM/1% XG were completely degraded and dissolved whereas structures printed using solely HA-PEG/1% XG remained intact. Structures comprising a combination of non-degradable and degradable features retained the latter but not the former unless the degradable hydrogel was completely covered by the non-degradable hydrogel, which resulted in trapping of the released HA and XG under the intact hydrogel. The slow diffusion of released HA through an intact hydrogel was further verified by casting a three-layer hydrogel structure in a cuvette. The bottom and top layers comprised of Cy3-labeled HA-PEG/1% XG hydrogels (pink) and the middle layer comprised of Cy5-labeled HA-VPM/1% XG hydrogel (blue) (Fig. 4c). Addition of Col-1 (0.5 mg ml^-1^) resulted in rapid degradation and substantial swelling of the middle hydrogel layer whereas the bottom and top layers appeared intact. Mechanical puncture of the intact top layer resulted in a release of the trapped degradation products and a collapse of the structure. Obviously, Col-1 was small enough to diffuse freely in the hydrogels whereas the degraded hydrogel components were not.

**Figure 4.**
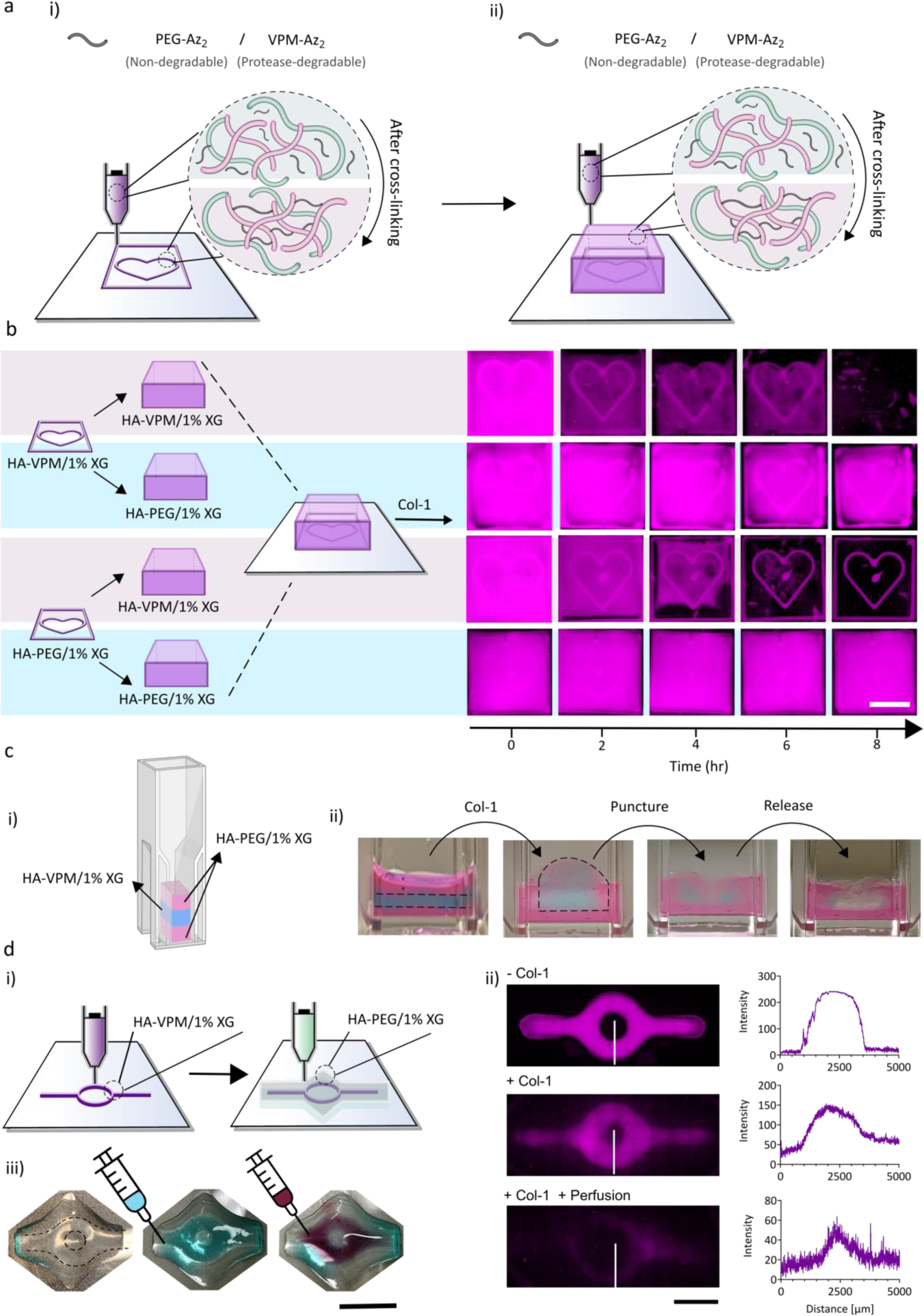
Protease remodeling of 3D bioprinted structures. **a.** Schematic illustration of the printing process and hydrogel composition. **b.** Proteolytic remodeling of printed structures by Col-1 using HA-VPM/1% XG and HA-PEG/1% XG hydrogels. Scale bar: 5 mm. **c.** Subjecting a layered structure comprising HA-PEG/1% XG (pink) and HA-VPM/1% XG (blue) to Col-1 resulted in degradation and swelling of the HA-VPM compartment. Physical disruption of the HA-PEG hydrogel released the degraded hydrogel components trapped under the intact top HA-PEG layer. **d.** i) Printing of HA-PEG/1% XG channel structures using Cy5-HA-VPM/1% XG hydrogels as a proteolytically degradable sacrificial bioink. ii) Fluorescence images (left) and intensity (right) of the channel before and after the addition of Col-1. The fluorescence intensity of the channel was monitored across the channel as indicated by the white line. iii) After incubation with Col-1, the channels allowed for free flow of buffer (blue/purple). Scale bars: 5 mm.

To further investigate the possibilities to use proteolytic activity for remodeling bioprinted structures, we utilized the HA-VPM/1% XG hydrogels as a sacrificial or templating ink to produce tubular structures inside an HA-PEG hydrogel. A channel path, 0.5 mm in width and 0.25 mm in height, was first printed using HA-VPM/1% XG and then covered by a printed layer of HA-PEG/1% XG (Fig. 4d). The final structures were subjected to 1 mg ml^-1^ Col-1 for 24 hours at 37 °C before the respective ends of the resulting channel were opened to allow for the degradation products to be released. The channels supported the free flow of the buffer and were mechanically stable. Given the general difficulties to create defined voids in soft materials using bioprinting, protease-responsive sacrificial bioinks can consequently provide new means for generating structural support when creating cavities or tubular structures that require cytocompatible materials and conditions for processing.

### Fibroblast and breast cancer cell mediated remodeling of 3D bioprinted structures

To further investigate the impact of proteolytic remodeling of the bioprinted structures by endogenous proteases secreted by the human breast cancer cells (MCF-7) and human primary fibroblasts, we created spatially defined tumor-mimicking microenvironments comprising different combinations of HA-VPM/1% XG and HA-PEG/1% XG. The two cell types were first investigated separately and then combined in the different bioprinted compartments. Structures comprising either HA-VPM/1% XG or HA-PEG/1% XG hydrogels embedded in fibroblast laden bioinks based on HA-VPM/1% XG or HA-PEG/1% XG were printed in the shape of a heart as illustrated in Fig. S4 (Supporting Information). Bioprinted fibroblast displayed an elongated morphology at the interface between the two compartments, but no fibroblasts were seen to penetrate the printed heart structures irrespectively of hydrogel composition (Fig. S5 and Fig. S6, Supporting Information). MCF-7 cells cultured in the bioprinted structures existed as individual cells or in small spheroids in all conditions for a period of 14 days (Fig. S7, Supporting Information). A significantly higher cell density was seen for MCF-7 cultured in HA-VPM/1% XG when covered by a printed layer of HA-VPM/1% XG compared to when cultured in HA-PEG/1% XG irrespectively of the covering hydrogel composition (Fig. S8, Supporting Information). The spheroid size was also larger when cultured in HA-VPM/1% XG and cells were on average more extended. No obvious proteolytic degradation of the heart structures was observed for any of the conditions (Fig. S4b-e, Fig. S5, Supporting Information). However, as described above, diffusion of degraded polymer components through layers of intact hydrogel is restricted. Proteolytic degradation of the hydrogels may thus contribute to both improved diffusion of oxygen and nutrients and possibilities for cell expansion and migration. Interestingly, dramatic differences in MCF-7 cell growth and spheroid size were seen when combining bioinks with MCF-7 and fibroblasts bioprinted in adjacent structures. (Fig. 5 and Fig. S9, Supporting Information). In all conditions, the fibroblasts promoted MCF-7 cell growth and the formation of MCF-7 spheroids, however, the MCF-7 cell density and MCF-7 spheroid volume were highly dependent on the composition and combination of hydrogels (Fig. 6b). Structures printed using solely HA-PEG/1% XG showed very few spheroids. Larger spheroids were obtained when using fibroblast-laden HA-VPM/1% XG bioinks but there was no significant increase in MCF-7 cell density. A distinct and significant increase in both MCF-7 growth and spheroid size was seen when printing the MCF-7 cells using HA-VPM/1% XG based bioinks. Interestingly, still there were no migration of fibroblasts into the printed MCF-7 heart structures in any of the conditions (Fig. S9, Supporting Information). However, MCF-7 bioprinted using HA-VPM/1% XG hydrogels and covered with bioprinted fibroblast laden HA-VPM/1% XG hydrogels showed tendencies to migrate out from their original compartment into the fibroblast compartment (Fig. 7a,b). For the other hydrogel compositions, the majority of the MCF-7 cells remined in their original compartment for the duration of the 14-day culture (Fig. 7c-f). MCF-7 cells that could escape the HA-PEG/1% XG and migrate into a fibroblast laden HA-VPM/1% XG compartment gained considerably in size (Fig. 7f). Hydrogel degradability in combination with the enhanced proteolytic remodeling of the hydrogels evoked by the fibroblast clearly facilitated MCF-7 migration towards the oxygen and nutrient gradient, in addition to promoting cancer cell proliferation. Interestingly, the compactness of the spheroids, defined as the number of nuclei divided by the cross-sectional area of the spheroids, varied significantly based on the hydrogel composition of the printed structures and the position of the spheroids (Fig. 7i,j). In general, spheroids in the heart structures were significantly more compact compared to spheroids located outside the heart structure irrespectively of the hydrogel compositions. This could indicate an accelerated growth of the cancer cells once they managed to migrate into the fibroblast compartment.

**Figure 5.**
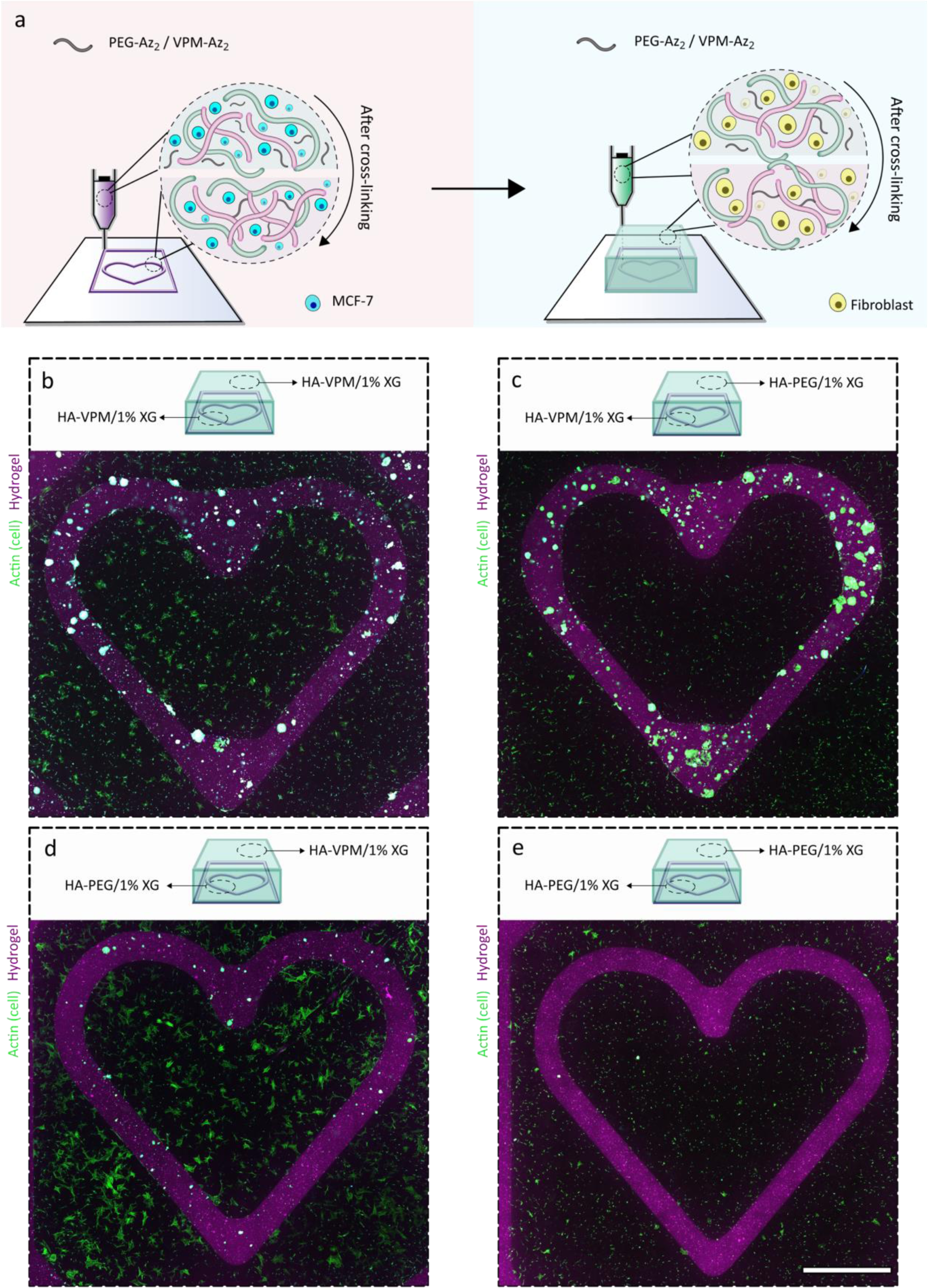
Bioprinting of MCF-7 and fibroblasts. **a.** Schematic illustration of the bioprinting process of structures containing MCF-7 (purple) covered by a homogenous hydrogel layer with fibroblast-laden bioink (green). **b.** Confocal micrograph of HA-VPM/1% XG structure (purple) containing MCF-7 covered by HA-VPM hydrogel containing fibroblasts. **c.** Confocal micrograph of HA-VPM/1% XG structure containing MCF-7 covered by HA-PEG/1% XG hydrogel containing fibroblasts **d.** Confocal micrograph of HA-PEG/1% XG structure containing MCF-7 covered by a HA-VPM/1% XG hydrogel with fibroblasts **e.** Confocal micrograph of HA-PEG/1% XG structure containing MCF-7 covered by HA-PEG/1% XG hydrogel with fibroblasts. Green: fibroblasts stained with phalloidin after 14 days of culture. Purple: Cy5-stained hydrogel. All images have been taken after 14 days of culture. Scale bar: 2.5 mm.

**Figure 6.**
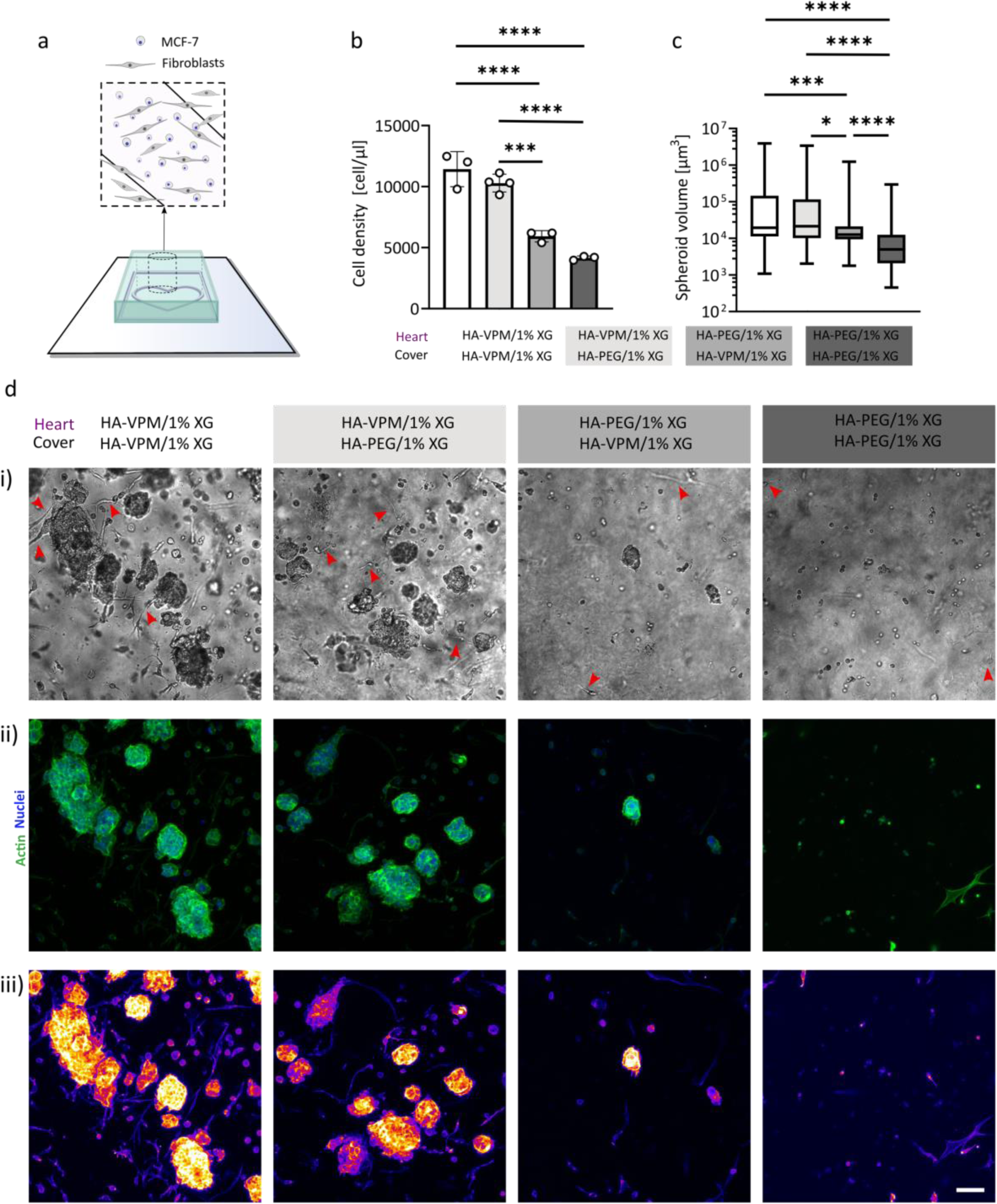
MCF-7 spheroid formation in bioprinted hydrogels. **a.** Schematic illustration indicating the part of the bioprinted structure that was imaged. The structure consisted of a heart containing MCF-7 and a hydrogel cover with fibroblasts. **b.** Quantification of MCF-7 cell density (****, P < 0.0001; ***, P < 0.0002, one-way ANOVA followed by Tukey’s multiple comparisons test, n=3). **c.** Quantification of MCF-7 spheroid size. (****, P < 0.0001; ***, P < 0.0002. Kruskal-Wallis test followed by Dunn’s multiple comparison test). **d.** Bright field (i) and fluorescence confocal (ii, iii) images of MCF-7 and fibroblasts after 14 days of culture. Red arrows in (i) indicate fibroblasts. In (ii) Green: phalloidin-stained F-actin, Blue: Hoechst-stained nuclei. In (iii), MCF-7 and fibroblasts are color coded orange/red and purple, respectively. Scale bar: 100 µm.

**Figure 7.**
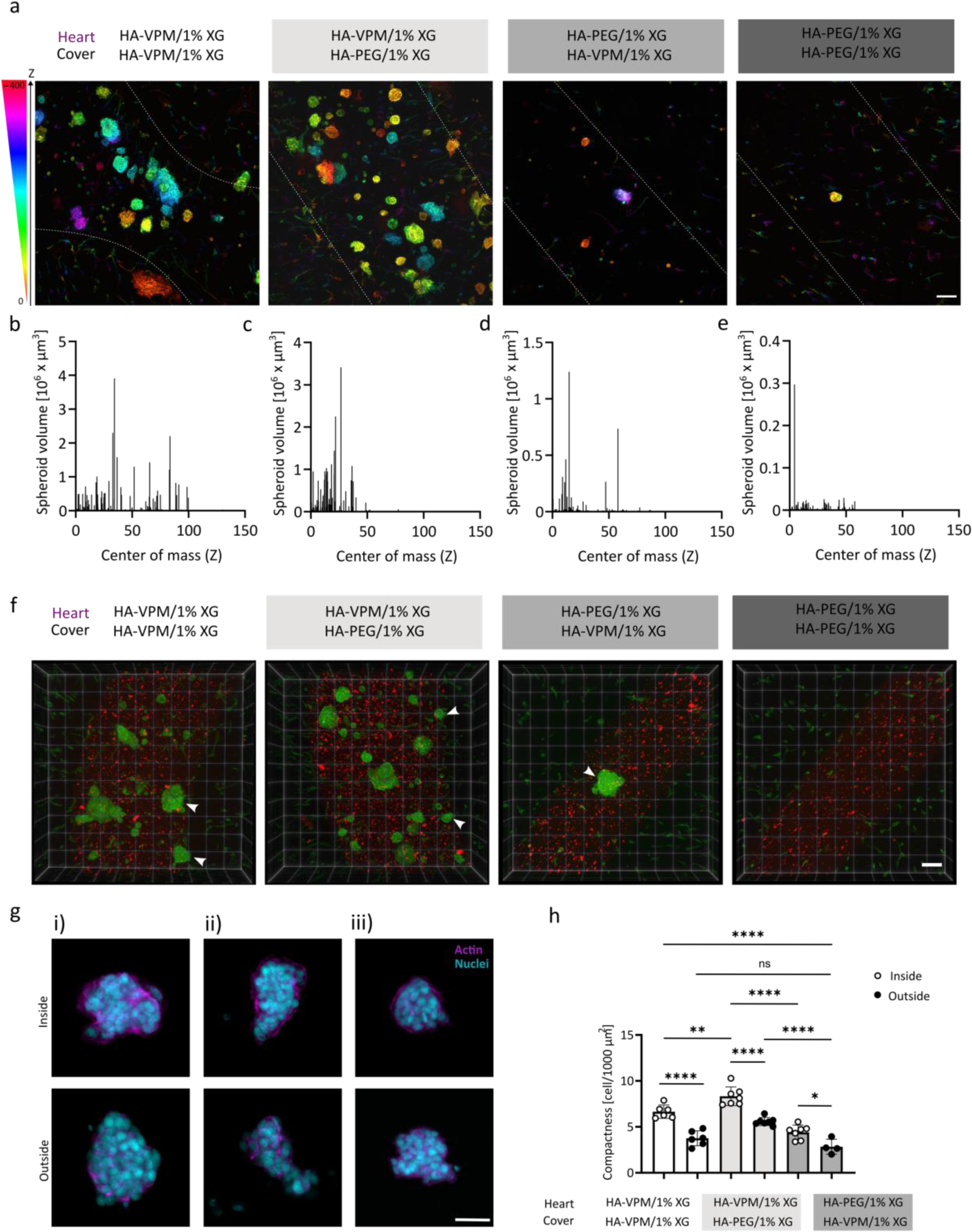
Analysis of MCF-7 spheroids in bioprinted hydrogels in contact with a fibroblast compartment. **a.** Color coded z-position of phalloidin stained MCF-7. Scale bar: 100 µm. MCF-7 spheroid size vs. spheroid center of mass along the z-axis in: **b.** HA-VPM/1% XG covered by HA-VPM/1% XG; **c.** HA-VPM/1% XG covered by a HA-PEG/1% XG; **d.** HA-PEG/1% XG covered by HA-VPM/1% XG; **e.** HA-PEG/1% XG covered by HA-PEG/1% XG. **f.** 3D confocal images of parts of the heart structures. White arrows indicate spheroids located outside the printed heart structure. Red: Cy5-stained heart, Green: phalloidin-stained F-actin. Scale bar: 100 µm. **g.** Representative confocal images of MCF-7 spheroids located inside and outside the bioprinted heart structure after 14 days of culture in: i) HA-VPM/1% XG covered by HA-VPM/1% XG; ii) HA-VPM/1% XG covered by HA-PEG/1% XG; iii) HA-PEG/1% XG covered by HA-VPM/1% XG. Magenta: phalloidin-stained F-actin, Blue: Hoechst-stained nuclei. Scale bar: 50 µm. **h**. compactness (number of cells per area of MCF-7 spheroid) outside and inside of the printed heart (****, P < 0.0001; ***, P < 0.0002; **, P < 0.0021; *, P < 0.0332. One-way ANOVA test with Tukey’s multiple comparison test.)

Cancer-associated fibroblasts (CAFs) can support and promote cancer growth, including breast cancer cells ^44, 45^, by expression and secretion of growth factors and various cytokines ^46^. Moreover, CAFs contribute to ECM remodeling by secretion of MMPs^47^. Taubenberger et al. have shown that MCF-7 spheroids became larger in protease degradable hydrogels compared to spheroids cultured in non-degradable hydrogels.^48^ Fibroblasts have also been seen to increase the invasiveness of breast cancer cell lines,^49^ potentially as a result of fibroblast contribution to upregulation and expression of MMPs from the MCF-7 cells ^50^. Our data aligns with these findings and clearly shows that the presence of fibroblasts in 3D bioprinted structure increased spheroid growth and cancer cell migration. These effects were significantly more pronounced when the hydrogels could be proteolytically remodeled.

As demonstrated here, the possibilities to combine and integrate bioinks with different susceptibilities to proteolytic remodeling allow for both new means for both protease-assisted biofabrication and fundamental studies of the multifaceted roles of proteases in tumor development and progression.

### Conclusions

In conclusion, we have developed materials and methods for 3D bioprinting of integrated hydrogel structures with selective response to proteolytic activity. The protease degradability of the hydrogels was tuned by altering the type of cross-linker, which was based on either a linear peptide (VPM) or PEG. Both hydrogels were functional as components in bioinks and supported high cell viability before and after printing and allowed for bioprinting with high lateral resolution when combined with xanthan gum (XG) to optimize the viscosity of the bioinks. The selective degradation of VPM-based hydrogels allowed for protease-assisted remodeling of bioprinted structures. The addition of collagenase type 1 (Col-1) resulted in the complete degradation of bioprinted features comprising VPM-cross-linked hydrogels but not hydrogels cross-linked using PEG. Bioprinting of microenvironments with different susceptibilities to proteolytic remodeling enabled a detailed investigation of fibroblast stimulated protease activity on MCF-7 proliferation and spheroid formation. Whereas limited effect of protease degradability was seen for MCF-7 alone, MCF-7 cultured in protease degradable hydrogels showed a significant increase in both proliferation and spheroid number and size in the presence of fibroblasts. When using protease degradable bioinks, the cancer cells migrated from their original bioprinted compartment into the fibroblast compartment resulting in formation of large spheroids. These findings demonstrate the central role of protease-mediated ECM remodeling on cancer cell migration and proliferation and highlight the complex interplay between tumor cells and fibroblast in this process. The possibility to design bioprinted structures with defined responses to proteolytic activity can facilitate both fabrication of dynamic bioprinted structures and complex tumor models for investigations of the crosstalk between cancer cells and the tumor microenvironment.

## Acknowledgments

Human dermal fibroblasts were a kind gift from Dr. Johan Junker (Disaster Medicine and Traumatology (KMC), Linköping University). Microscopy core facility at Linköping University. The financial support from the Knut and Alice Wallenberg Foundation (KAW 2016.0231, 2021.0186) and Carl Tryggers Stiftelse is gratefully acknowledged.

## Notes

### Competing Interest Statement

The authors have declared no competing interest.

