## Supplementary information for "Proteolytic Remodeling of 3D Bioprinted Tumor Microenvironments"

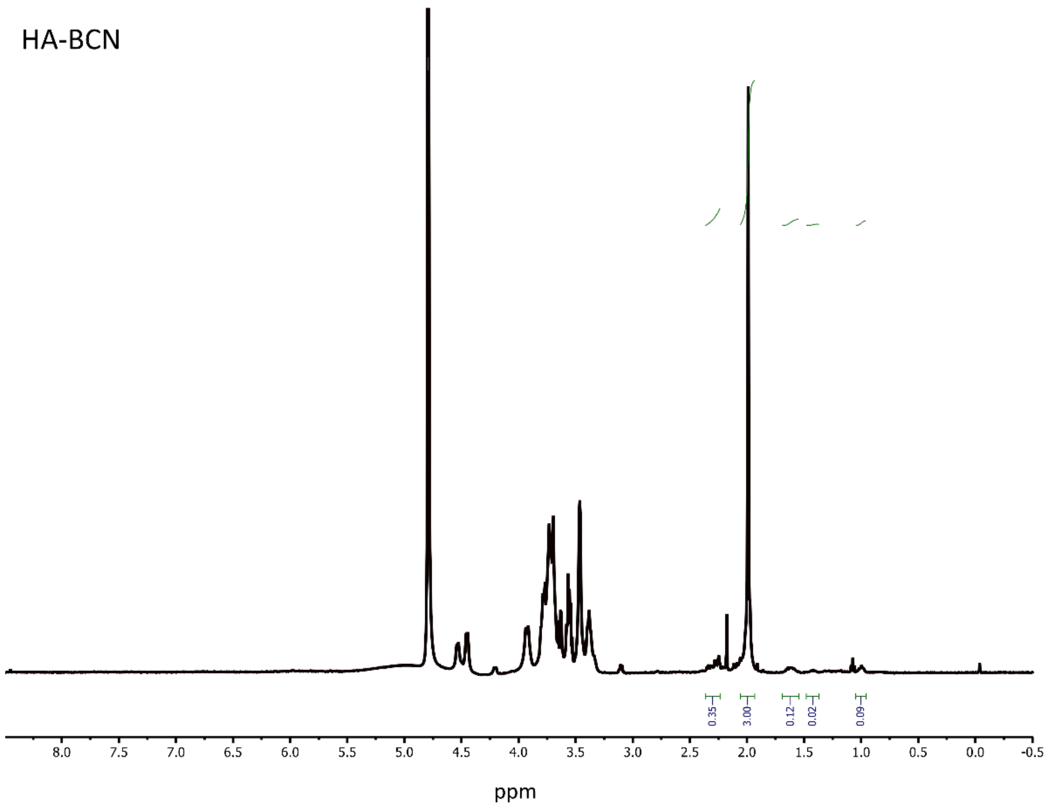

**Figure S1.** [1]H-NMR spectra of HA-BCN in $D_2O$. Peaks used to calculate the degree of functionalization of HA with BCN groups are marked. A degree of functionalization of 7% was calculated based on marked integrals and normalized with respect to the peak deriving form *N*-acetyl (3H) in HA at 1.95 ppm.

VPM peptide sequence

Ac- Lys($N_3$) - Gly - Arg - Asp - Val - Pro - Met - Ser - Met - Arg - Gly - Gly - Asp - Arg - Lys($N_3$) -CONH2

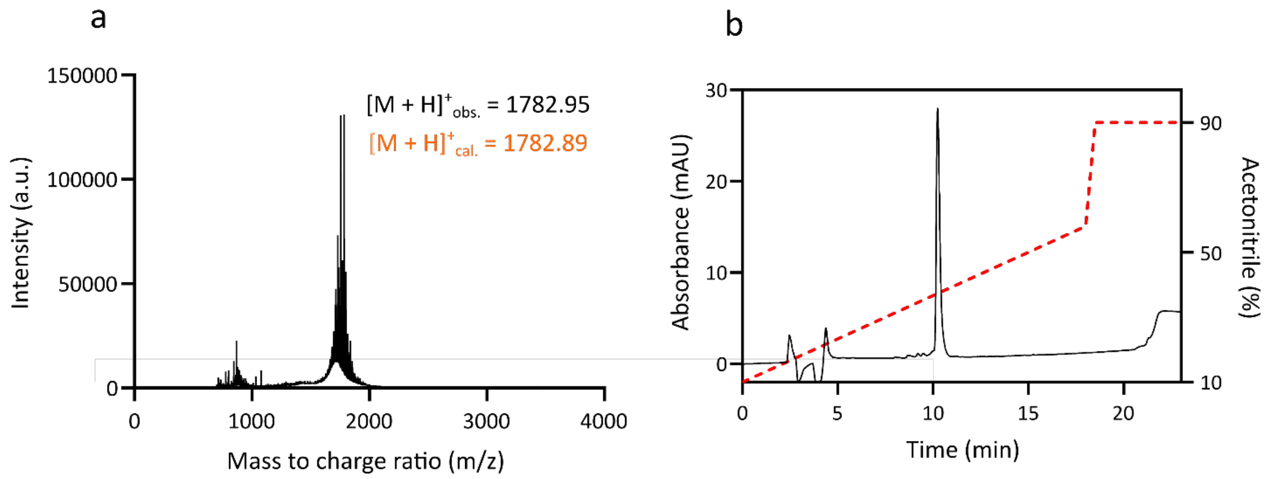

**Figure S2:** Characterization of VPM-$Az_2$ after purification. **a.** Mass spectra of VPM-$Az_2$. **b.** HPLC trace of VPM-$Az_2$ in a gradient of aqueous acetonitrile ranging from 10% to 90% with 0.1% TFA added.

**Table S1.** Optimization of hydrogel composition.

| Volume of VPM-$Az_2$ (20 mg $ml^{-1}$) / PEG-$Az_2$ (12.34 mg $ml^{-1}$) | The volume of HA-BCN (20 mg $ml^{-1}$) | Mechanical properties* |
|---|---|---|
| 0 | 20 | Negative control |
| 0.5 | 19.5 | + |
| 1 | 19 | ++ |
| 2.5 | 17.5 | +++ |
| 5 | 15 | ++ |
| 10 | 10 | + |

* + very soft, ++ soft, +++ acceptable stiffness

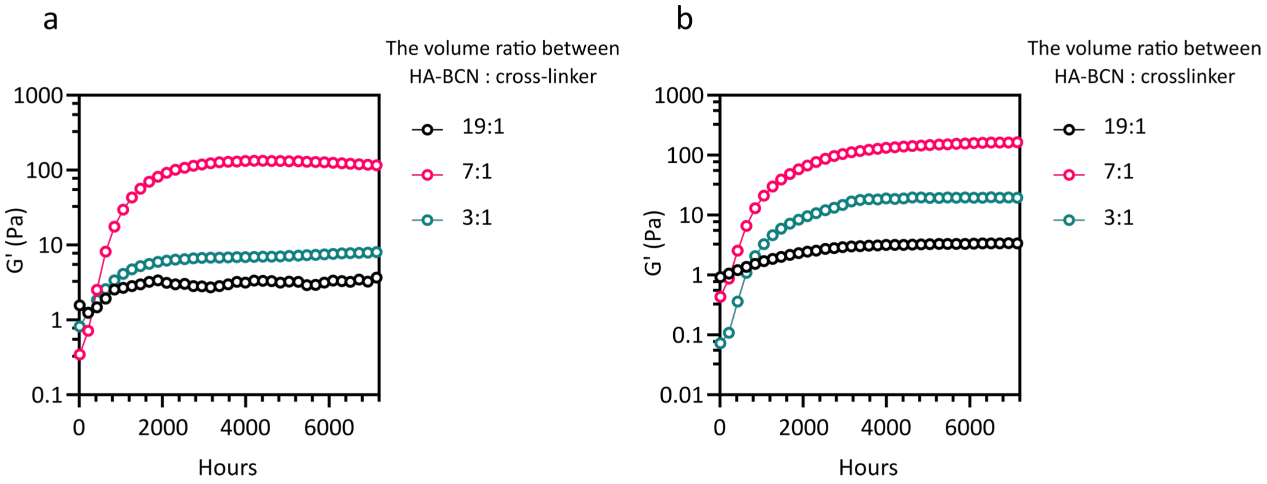

**Figure S3**. Hydrogel cross-linking kinetics for different hydrogel compositions. **a.** HA-BCN : VPM-$Az_2$ and **b.** HA-BCN : PEG-$Az_2$.

**Table S2.** Bioprinting parameter for different hydrogel compositions.

| | Hydrogel composition | | Printer settings | | | |
|---|---|---|---|---|---|---|
| Name | HA-BCN % | Xanthan Gum % | Print bed temp. | Nozzle size | Speed mm/s | Pressure kPa |
| HA | 2 | 0 | RT | 25G | 2-4 | 0-1 |
| HA-0.5 % XG | 2 | 0.5 | RT | 25G | 2-4 | 0-1 |
| HA-1% XG | 2 | 1 | RT | 25G | 1-2 | 3-6 |
| HA-1.5% XG | 2 | 1.5 | RT | 25G | 1-2 | 5-11 |

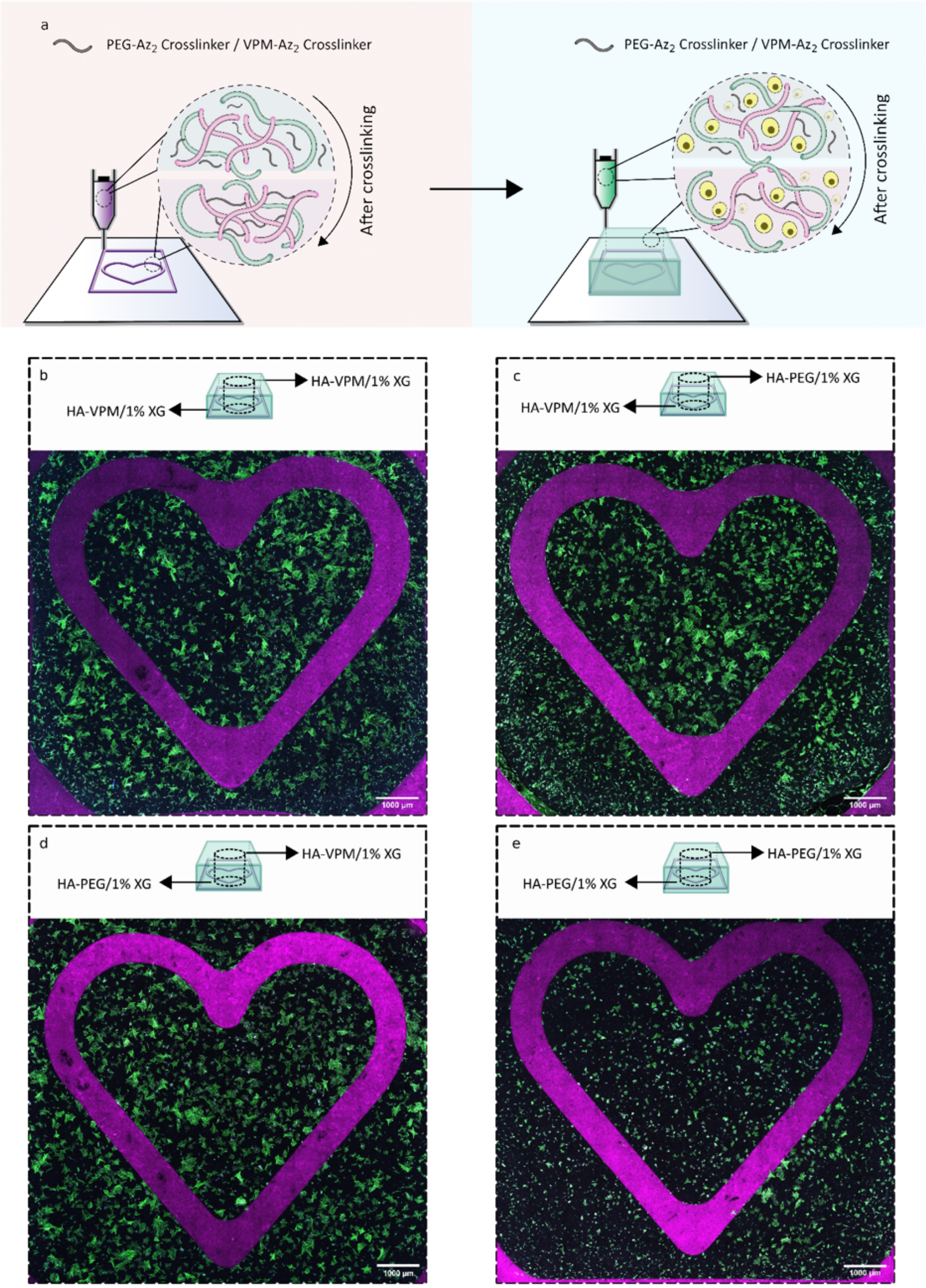

**Figure S4.** 3D bioprinting of fibroblasts. **a.** Schematic illustration of the bioprinting process of non-cell containing structures (heart, purple) and cell-laden hydrogels containing fibroblasts. Two different types of bioinks were used to print the structures, using protease degradable (VPM-Az$_2$) and non-degradable (PEG-Az$_2$) crosslinkers and 1% XG. **b-d.** Confocal images of 3D bioprinted structures (purple, Cy5-stained hydrogel) and covered by a bioprinted hydrogel containing fibroblasts (green, phalloidin) after 14 days of culture using hydrogel compositions as indicated in the corresponding schematic.

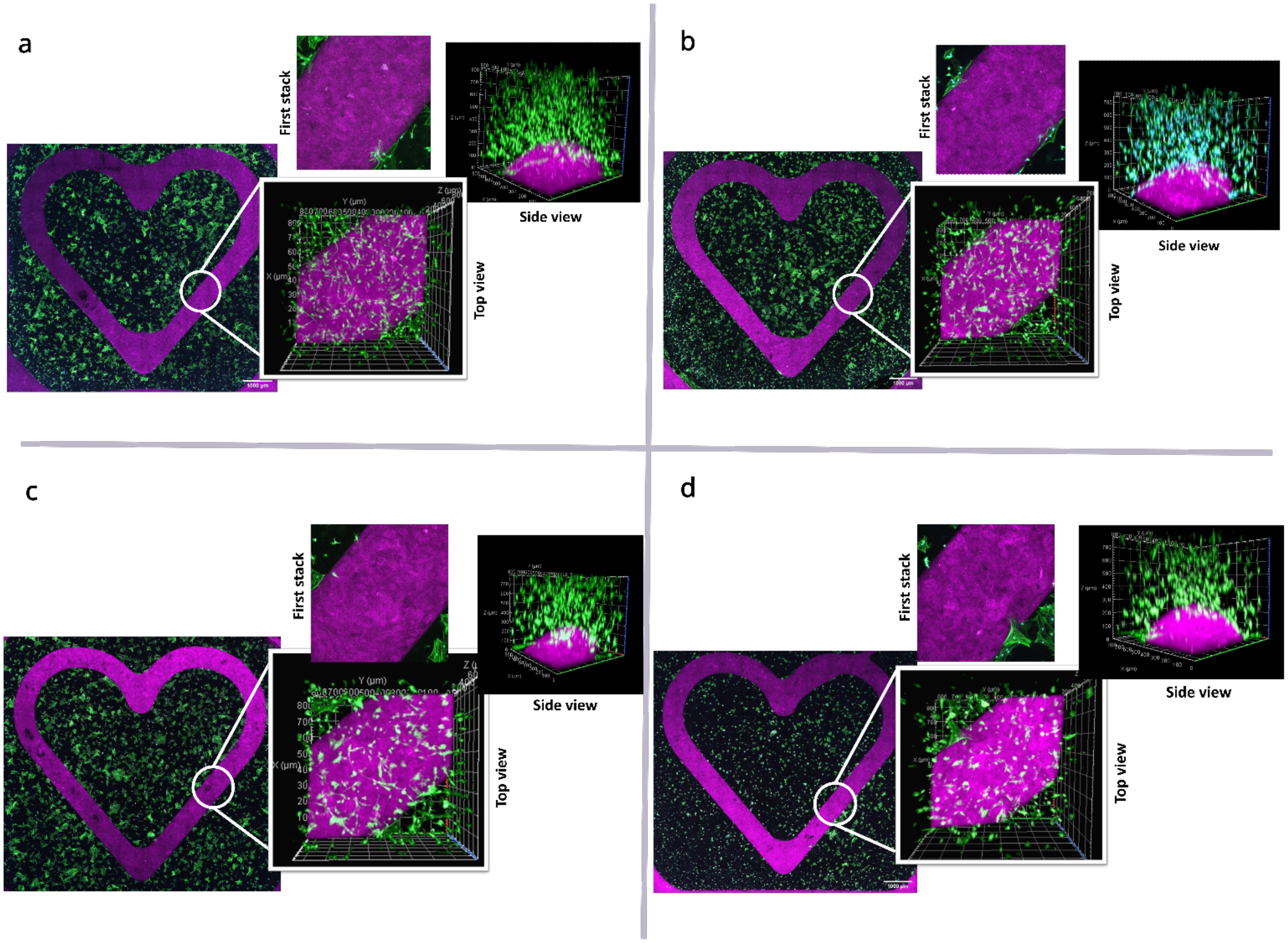

**Figure S5.** Confocal images of 3D bioprinted structure after 14 days of culture. **a.** Bioprinted structure of HA-VPM/1% XG covered with fibroblasts containing HA-VPM/1% XG. **b.** Bioprinted structure of HA-VPM/1% XG covered with fibroblasts containing HA-PEG/1% XG. **c.** Bioprinted structure of HA-PEG/1% XG covered with fibroblasts containing HA-VPM/1% XG. **d.** Bioprinted structure of HA-PEG/1% XG covered with fibroblasts containing HA-PEG/1% XG.

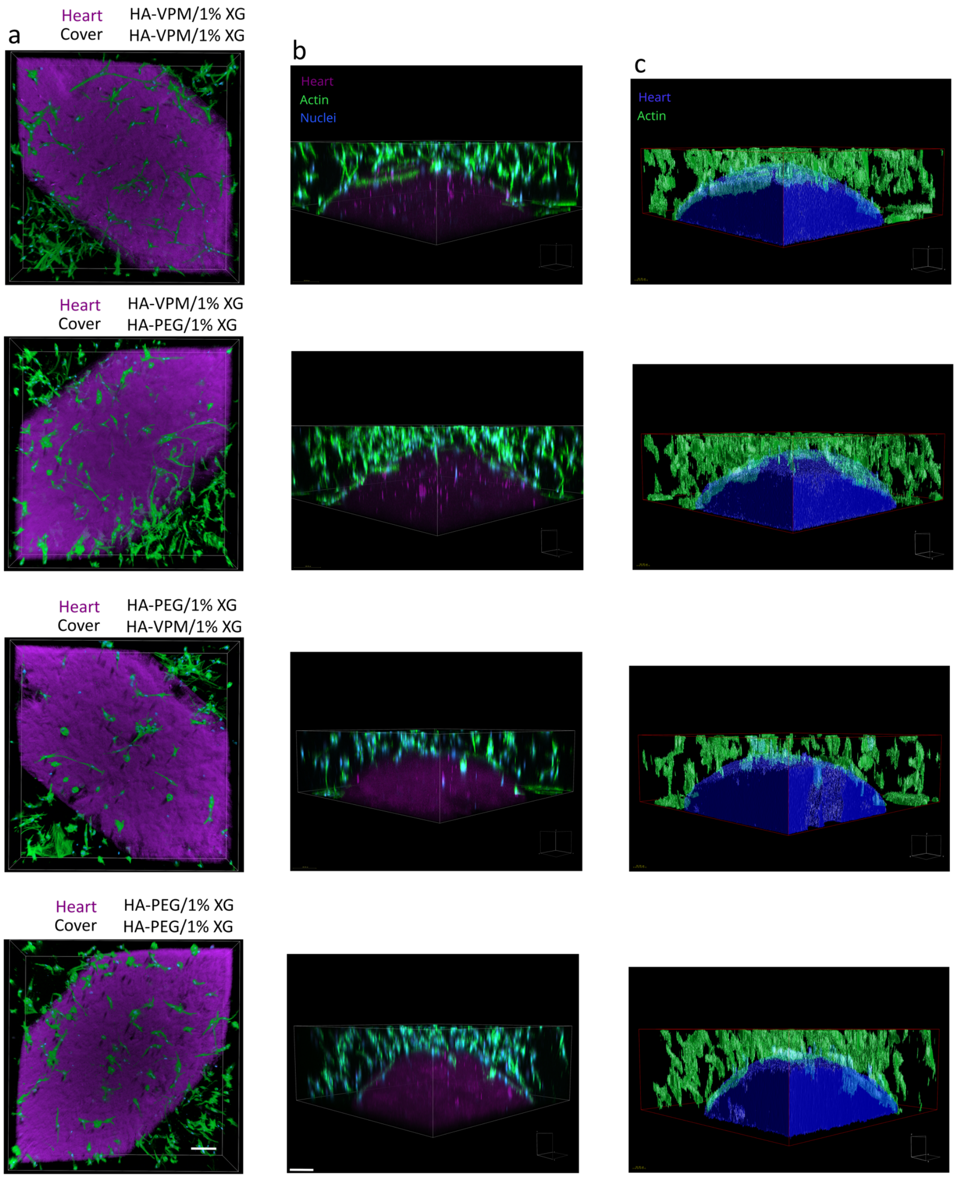

**Figure S6.** Images of 3D bioprinted structure containing fibroblasts after 14 days of culture. **a.** SFP (the Simulated Fluorescence Process) rendered images of bioprinted structures. **b.** Confocal images of 3D bioprinted structures after 14 days of culture (side view) **c.** Surface rendered images of 3D bioprinted structure after 14 days of culture (side view). Scale bar: 100 µm.

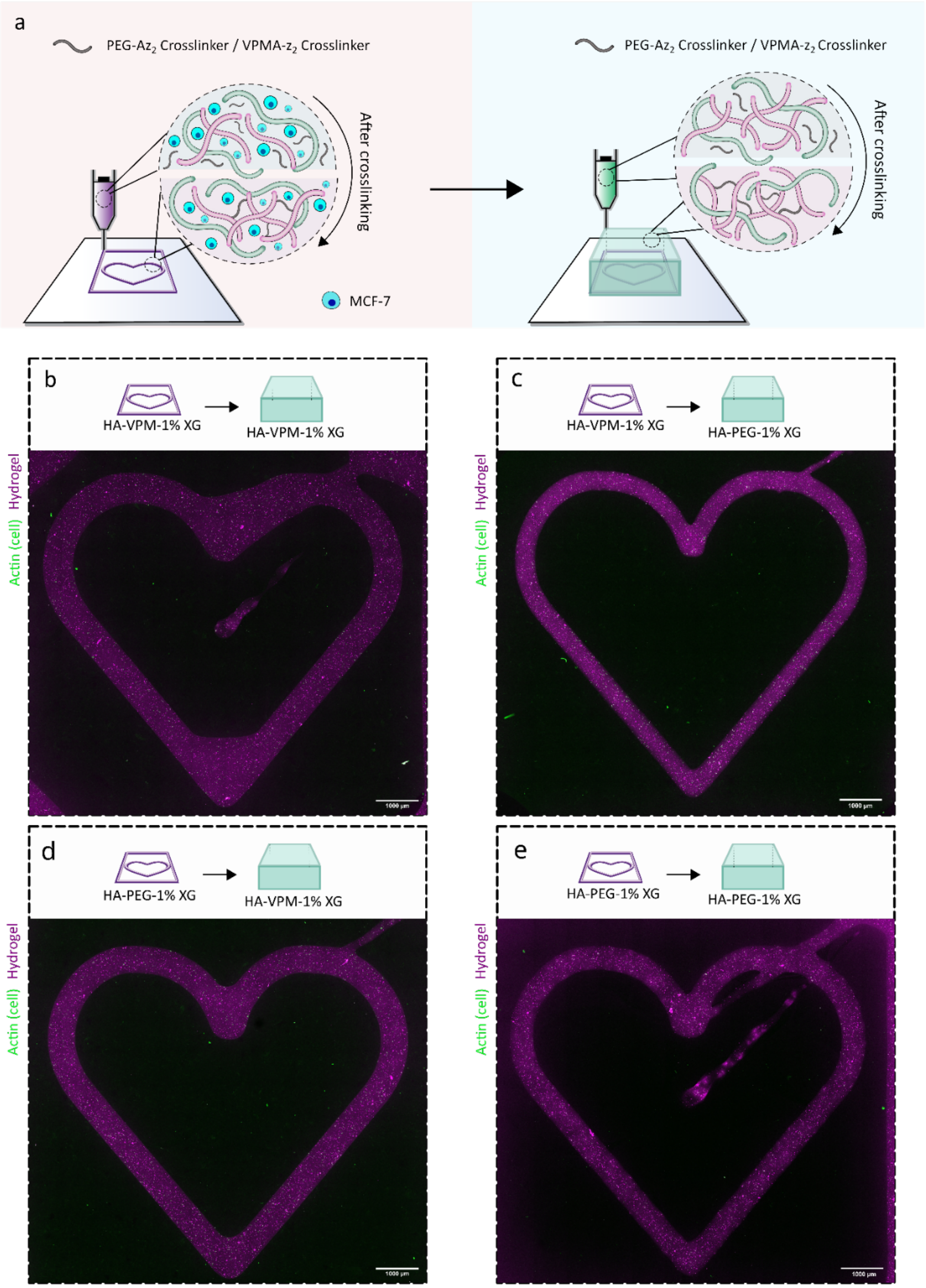

**Figure S7.** 3D bioprinting of MCF-7. **a.** Schematic illustration of the bioprinting process of cell laden hydrogel containing MCF-7 (heart, purple) and non-cell containing hydrogels. Two different types of bioinks were used to print the structures, using protease degradable (VPM-$Az_2$) and non-degradable (PEG-$Az_2$) cross-linkers and 1% XG. **b-d.** Confocal images of 3D bioprinted structures (purple, Cy5-stained hydrogel) containing MCF-7 (green, phalloidin) and covered by a bioprinted hydrogel after 14 days of culture using hydrogel compositions as indicated in the corresponding schematic.

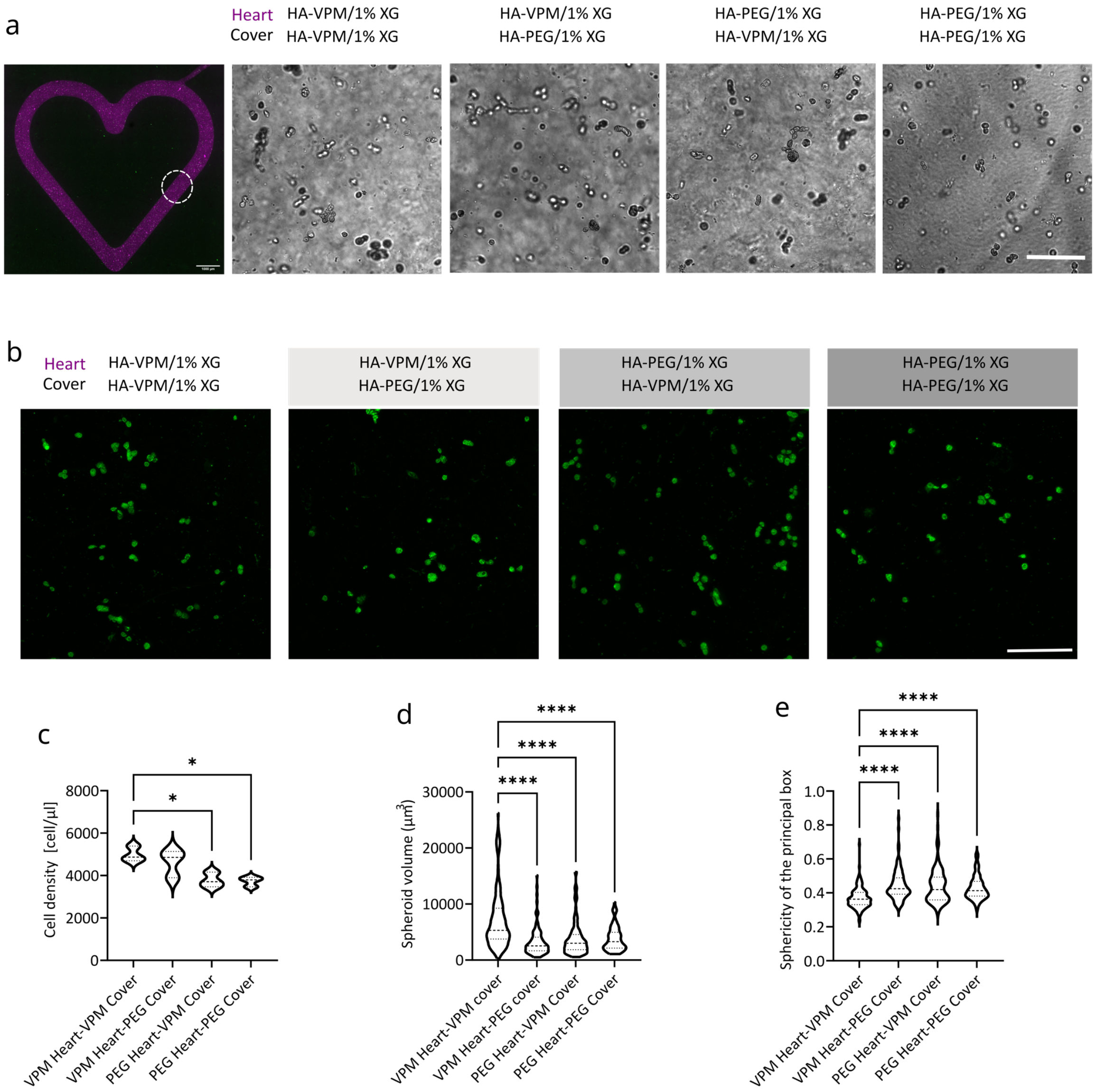

**Figure S8.** Images and analysis of 3D bioprinted structures containing MCF-7 after 14 days of culture. **a** Confocal image of bioprinted structure (purple, Cy5-stained hydrogel) containing MCF-7 covered with a printed hydrogel and corresponding bright field images for the different conditions, scale bar: 100 µm. **b.** Confocal images of MCF-7 (Green: phalloidin-stained F-actin) in bioprinted structures, scale bar: 100 µm **c.** Cell density, **d.** Spheroid size, **e.** Sphericity of MCF-7 cultured in the bioprinted structures.

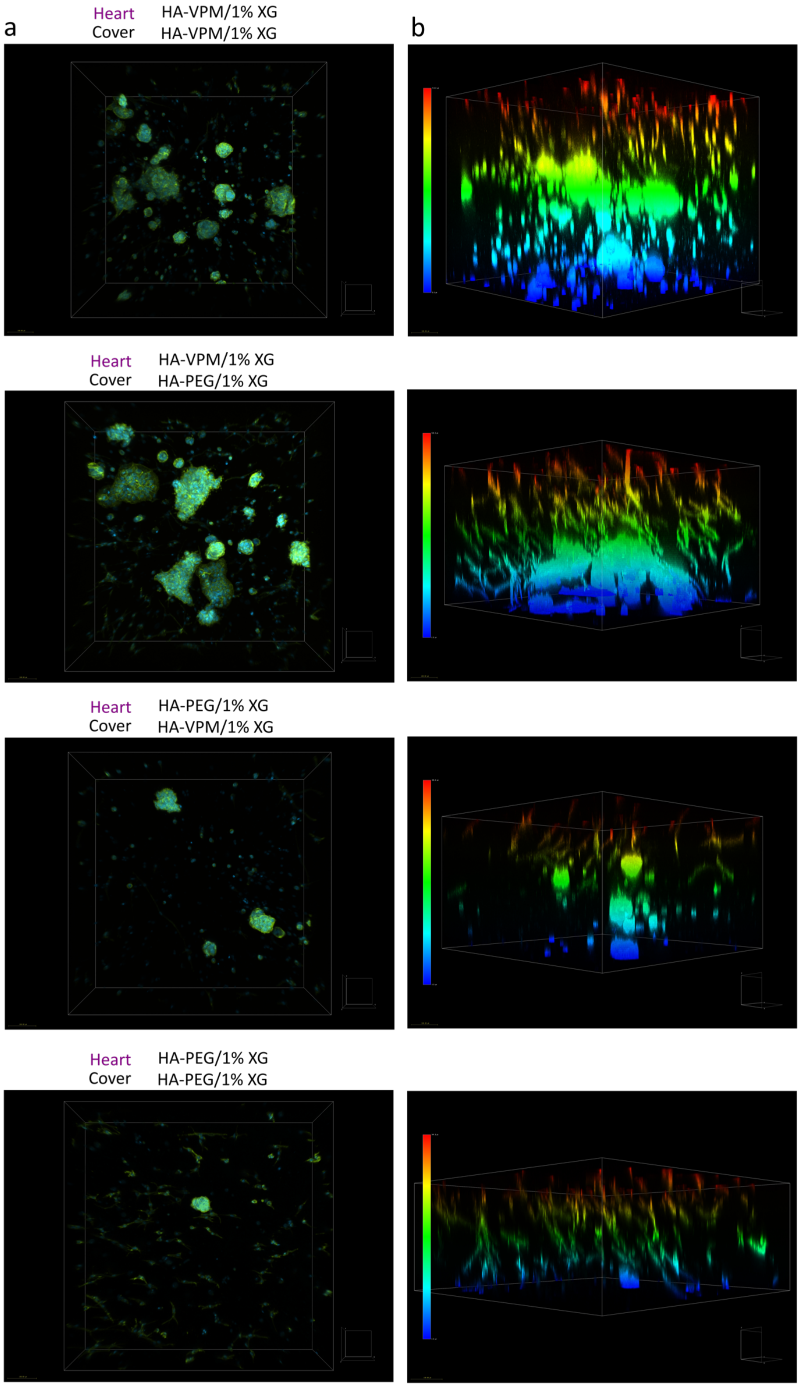

**Figure S9.** Images of MCF-7 and fibroblasts in 3D bioprinted structures after 14 days of culture showing no migration of fibroblasts into the heart structures. **a.** 3D confocal images of MCF-7 and fibroblasts in 3D bioprinted structure after 14 days of culture. Green: phalloidin-stained F-actin, Blue: Hoechst-stained nuclei. **b.** A spectrum LUT was applied to representative confocal images to indicate the height of MCF-7 and fibroblasts in the structure.
